# Development of LSTM&CNN Based Hybrid Deep Learning Model to Classify Motor Imagery Tasks

**DOI:** 10.1101/2020.09.20.305300

**Authors:** Caglar Uyulan

## Abstract

Recent studies underline the contribution of brain-computer interface (BCI) applications to the enhancement process of the life quality of physically impaired subjects. In this context, to design an effective stroke rehabilitation or assistance system, the classification of motor imagery (MI) tasks are performed through deep learning (DL) algorithms. Although the utilization of DL in the BCI field remains relatively premature as compared to the fields related to natural language processing, object detection, etc., DL has proven its effectiveness in carrying out this task. In this paper, a hybrid method, which fuses the one-dimensional convolutional neural network (1D CNN) with the long short-term memory (LSTM), was performed for classifying four different MI tasks, i.e. left hand, right hand, tongue, and feet movements. The time representation of MI tasks is extracted through the hybrid deep learning model training after principal component analysis (PCA)-based artefact removal process. The performance criteria given in the BCI Competition IV dataset A are estimated. 10-folded Cross-validation (CV) results show that the proposed method outperforms in classifying electroencephalogram (EEG)-electrooculogram (EOG) combined motor imagery tasks compared to the state of art methods and is robust against data variations. The CNN-LSTM classification model reached 95.62 % (±1.2290742) accuracy and 0.9462 (±0.01216265) kappa value for datasets with four MI-based class validated using 10-fold CV. Also, the receiver operator characteristic (ROC) curve, the area under the ROC curve (AUC) score, and confusion matrix are evaluated for further interpretations.

## 1. Introduction

The use of assistive technologies to help disabled people is importantly increasing in recent years and researchers propose inspirational new scientific methods for restoring functions to those with motor impairments such as paralysis, amyotrophic lateral sclerosis, cerebral palsy, loss of limb [10]. Recent neuroscience and robotic studies indicate that even the imagination of a movement generates the same mental pattern as the performance of the movement itself [13]. Thus, transforming a brain activity or a task signal into direct control of any hardware device without the involvement of the peripheral nervous system or muscle is applicable. Therefore, it becomes promising for subjects suffering from mobility reduction. The techniques that enable the researchers to translate and interpret the brain signals as physical tasks are referred to as BCI [44]. BCI application provides a means of non-muscular communication and control paradigm to transmit signals&commands from the individuals with severely impaired movement to the external world or devices by measuring brain activity.

BCI technology is broadly composed of five consecutive processes, which are sequenced as; *signal acquisition (1), extraction of the intended action or desired features from the task (2), selection of more relevant subset from the feature set (3), classification of the mental state (4), and finally, feedback signals generated by the prosthetic device (5)* [38]. These brain signals are extracted, decoded and studied with the help of various imaging techniques like EEG, EOG, magnetoencephalography (MEG), positron emission tomography (PET), functional magnetic resonance imaging (fMRI), electrocorticography (ECoG), etc.[11].

But, since these techniques involve sophisticated and expensive equipment and therefore the availability and utilization are mostly assigned to high budget corporations or hospitals, the traditional measurement of brain activity, in the context of EEG-based BCI, has relied on the acquisition of EEG data via non-invasive electrode arrays. The acquired EEG data is analyzed in either the time or frequency domain dynamics and subsequently translated into corresponding control commands. A BCI technology aims at decoding characteristic brain modalities, which are generated from various brain locations, to control a robotic device. In the non-invasive method, the neurophysiological rhythms are recorded by sensors placed over the scalp to avoid the risks of surgery with a trade-off of low signal-to-noise (SNR) [22].

MI-related BCI, which is based on the imagination of the execution of movements is widely implemented. MI stimulates similar brain pathways as if it was performed a real movement. It replaces the exercises in cases where there is no residual motor function [48]. Several studies in the literature have focused on classifying MI-EEG signals to provide feedback during MI training [46; 65; 1]. While performing MI tasks, the synchronized rhythmic variations called event-related synchronization (ERS) and event-related desynchronization (ERD) in sensorimotor regions can be captured from EEG [24].

In the literature, several feature extraction and machine learning (ML)-based classification techniques are used for EEG-based BCI. Generally, feature extraction methods for the MI-EEG focus on deriving time-domain features, i.e. energy and amplitude of the signal, autoregressive modelling [17; 2], and on establishing frequency domain features [31; 37] or on extracting time-frequency features [59; 58]. With its adaptive structure and the ability for analyzing the non-stationary signals wavelet transform (WT) keeps and processes both time and frequency components of the signal. The combination of useful frequency and time information on the non-stationary EEG signal improves the performance of classifiers. Stating the merit of potential biomarkers is a critical threshold contributing to the classification performance. Therefore, with the proliferation of high-dimensional data, feature selection (FS) methods have been widely applied as a vital task before the learning process. The purpose of using FS methods is selecting a valuable subset of features from the original set of features without sacrificing from the accuracy in representing the initial set of features, in which plenty of spurious information and irrelevant features exist [52]. Extensive implementation of wrapper-based approaches is also underlined, particularly in bioinformatics, employing genetic algorithm (GA) [35], particle swarm optimization (PSO) [47], ant colony optimization (ACO) [16], etc. Besides, an increasing number of researches make use of the embedded capacity of several classifiers to discard less informative input features. ML-supported classification techniques such as quadratic discriminant analysis (QDA), linear vector quantization (LVQ), k-nearest neighbour (KNN), multilayer perception (MLP), support vector machine (SVM), linear discriminant analysis (LDA), decision tree, naive Bayesian classifier for EEG-MI classification have been widely studied [18; 14; 54; 62]. Since the brain signals recorded using EEG have non-linear, complex, non-stationary and non-Gaussian nature, finding a robust and accurate feature extraction and ML-based classification method is a challenge in EEG-based BCI application. To improve the effectiveness of the classifiers, the specialized pre-processing, artefact removal, feature reduction techniques can be used [23; 26; 63]. PCA is one of these techniques. With the aid of PCA, the higher dimensions of the signal, which contains relatively insignificant or insensitive features, are reduced to lower dimensions to increase the correctly classified percentage of data. It tries to find the “best” eigenvalues in the sense of variances while accounting the temporal variability. This means, that it is sought to extract the most dynamic one, but this does not necessarily mean that these features are the most prominent ones. Therefore DL algorithms should be applied for solving this problem.

DL is a prediction method, which uses a sequence of nonlinear processing stages, which jointly learns from data. It serves a new way of neural network (NN) fitting approach with hierarchical feature extraction and helps to find the representations that are invariant to inter-and intra-subject differences while reducing dimensions, in this way it is possible to construct a unified end-to-end model that can be applied to raw signals [4].

Deep NN’s have specialized and proved their effectiveness in recognition tasks including applications of images, videos, speech, and text classification. CNN’s are very suitable to study with images and video data because they are capable to extract representative features, which are robust to partial translation and deformation of inputs [30; 5]. CNN’s are also effective in many applications, which comprise temporal dynamics such as, handwriting, speech recognition. Additionally, CNN’s are utilized in the field of the combination of spatial representation and time-series structure, i.e. moving object detection or video classification [40; 33]. CNN’s provide significant performance enhancement minimizing the error rates of competing techniques in ImageNet competition 2012 [28].

In this paper, a continuous classification output for each sample in the form of class labels of MI tasks (0 (Left Hand), 1 (Right Hand), 2 (Feet), 3 (Tongue)) including labelled trials was provided by implementing CNN-LSTM based deep classifier based on the BCI Competition IV dataset A. The classification algorithm is causal, meaning that the classification output at time *k* may only depend on the current and past samples *x*_*k*_, *x*_*k*-1_, …, *x*_0_. A confusion matrix was then built from all artefact-free trials for each time point. The time course of the accuracy and loss was obtained. The mean and standard values of the validation accuracy and validation kappa value after 10-fold CV are computed, respectively. The confusion matrix and ROC curve were plotted, and the AUC score is evaluated.

### 1.1. Related studies

A unified end-to-end CNN-based DL model was developed to classify MI-related tasks. Transfer learning was used to adapt the global classifier to single individuals improving the overall mean accuracy. However, in this study, the classification performance for four class is quite low with 68.51% [15]. A novel approach for learning deep representations from multi-channel EEG time-series was proposed and its advantages in the mental load classification conceptualization were demonstrated. A deep recurrent-convolutional network was trained by mimicking the video classification techniques. As a result, the spatial, spectral, and temporal dynamics of EEG were preserved and mental load classification performance and robustness were improved [5]. A tensor-based multiclass multimodal scheme for hybrid BCI was developed to generate nonredundant tensor components. Multimodal discriminative patterns were selected through a weighted fisher criterion and support vector machine (SVM) was used for multiclass classification. The main advantage of this method is to capture the interactive effects of simultaneous tasks, but tensor generation and decomposition processes are very time-consuming [25]. Deep ConvNets with a range of various architectures including batch normalizations, exponential linear units, cropped training strategy were designed for decoding imagined or executed movements acquired from raw EEG. This method was compared with a validated baseline method named as filter bank common spatial patterns (FBCSP) decoding algorithm. One of the important findings of this study is that the deep ConvNets are learned features different from FBCSP, which could explain their higher accuracies in the lower frequencies where band power may be less important. Deep ConvNets can learn band power features with specific spatial distributions from the raw input in an end-to-end manner [51]. A novel MI classification framework was introduced by building a new 3D representation of EEG and training a multi-branch 3D CNN. Experimental evaluations reveal that the framework reaches state-of-the-art kappa value and outperforms other algorithms by a 50% decrease in the standard deviation of various subjects in terms of robustness criteria [66]. A CNN was employed to classify and characterize the error-related brain response as measured in 24 intracranial EEG recordings. It was found that the decoding accuracies of CNNs were higher than those of regularized linear discriminant analysis. The 4-layered CNNs were able to learn in all-channel decoding of errors from intracranial EEG electrodes in epilepsy patients [60]. It was proved that the CNN and LSTM capacity to learn high-level EEG features consisted of low-level ones, after feature extraction by discrete wavelet transform (DWT). The CNN and LSTM schemes are suitable and relatively roust to the BCIs and MI-EEG decoding [64].

## 2. THEORETICAL BACKGROUND

### 2.1. CONVOLUTIONAL NEURAL NETWORK

CNN’s are a kind of NN having a grid-like topology, which is specialized for processing data. The CNN’s are capable of extracting spatial features to a granular level and these features perform a high discrimination power for classification issues. A typical CNN architecture consists of three layers: convolution, pooling, and fully connected. Each layer comprises filter windows that slide over the input layer from the preceding layer.

While time-series data represents a 1-D grid, image data express 2-D grid nature comprising pixels. They take their name from a mathematical operation defined as *“convolution operator”*. CNN’s use the convolution operator instead of the general matrix multiplication in at least one of its layers [21]. The convolution of two discrete signals *x*_*n*_ and *ω*_*n*_ is given as 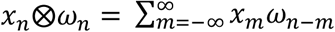, where ⨂ corresponds to the convolution operator. The generalized architecture of CNN is demonstrated in Fig.1.

**Figure 1:**
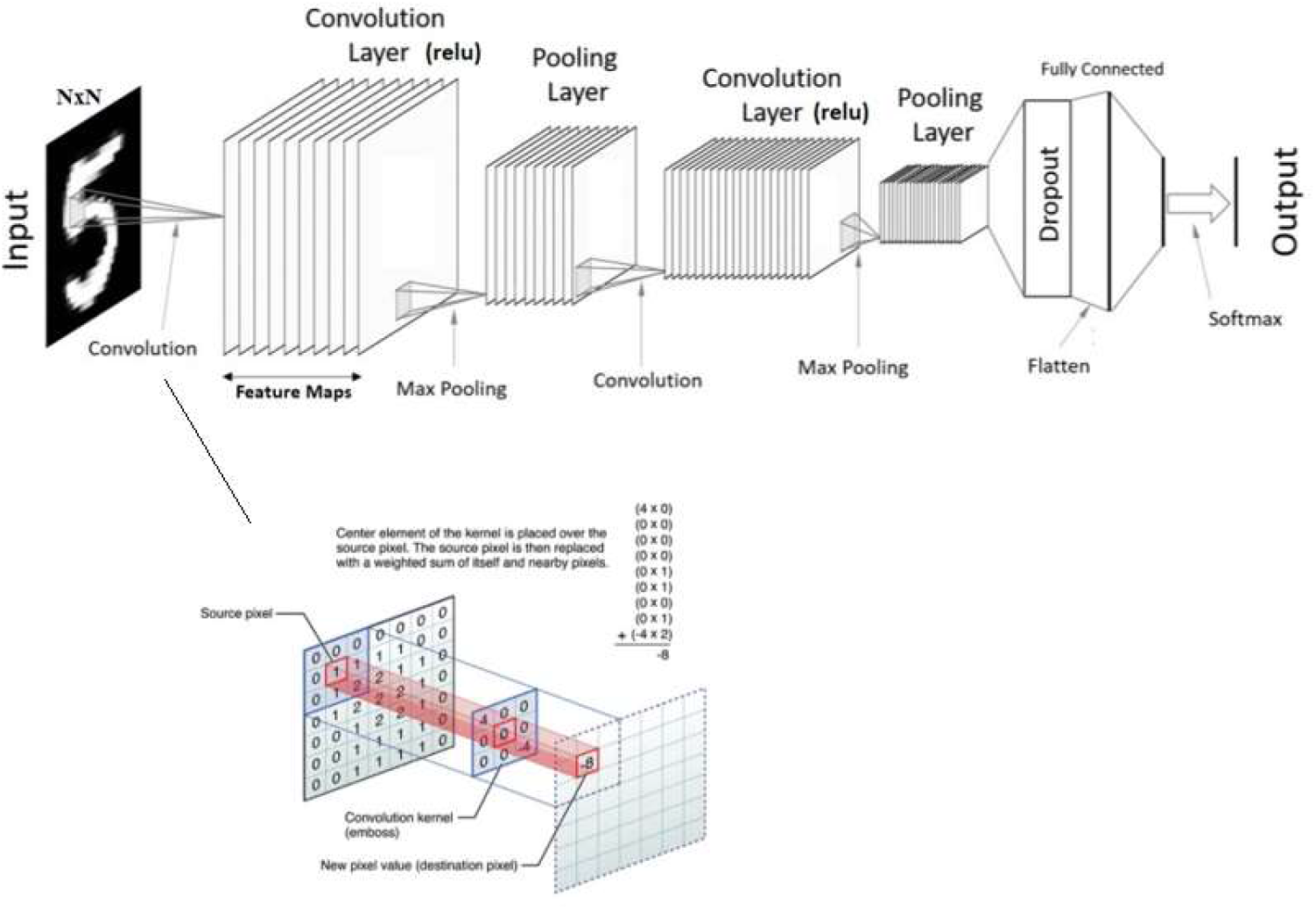
The generalized architecture of CNN’s.

Each neuron in the first layer of the CNN interacts only with a small region of the input neurons, which is defined as a convolution window (Fig.1). While the convolution window is passing through an entire input sequence, each neuron in the hidden layer learns by changing its connection weight and overall bias. The size of the convolutional window is named as the *“kernel size, k”*. The mathematical expression of this process is given as in Eq.1

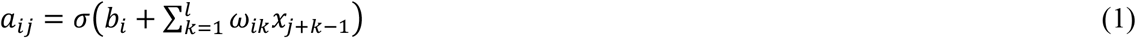

where *a*_*ij*_ is the output of the *j*^*th*^ neuron of the *i*^*th*^ filter in the hidden layer, *b*_*i*_ denotes to the overall bias of filter *i, ω* corresponds to the shared weights and *σ*(.) is the nonlinear activation function.

It can be deduced that if it is given a finite kernel size (*k*), the input to a specific neuron only relies on the subspace from the previous layer. This implies the sparse connectivity [43]. The sparse connections and weight sharing attribute extremely reduce the number of weights to be learned and shorten the training process by decreasing the gradient computation process. In this way, complex and high-level features are possible to be learned while preventing overfitting. Multiple filtered forms of the input data take place in the stacked hidden layers of the CNN as feature maps. Filter size, strides (sliding of windows), and padding (window offset over input) settings are parameterized.

Another operation is named as *“pooling”*. Through the pooling operation, high-dimensional input space is gradually confined to a low dimensional space between layers by maximizing or averaging, etc its neighbouring values in the feature map. The location independence of the model is increased because the feature in various positions can be mapped to the same feature through the help of the aggregation of adjacent neurons [53]. *“Fully connected layer”* structure behaves same as ANN. Each neuron is connected to every other neuron of the preceding layer. The theoretical basics of the CNNs and learning theories can be investigated more detailed from [32; 8; 39].

### 2.2. LONG-SHORT TERM MEMORY

LSTM networks are the modified version of the recurrent neural networks (RNN). RNNs are specialized to process sequential data (*x*^(1)^, *x*^(2)^, …, *x*^(τ)^), which plays an over time such as speech, language, audio, video etc. RNNs can scale long sequences and also process sequences of variable length and context. RNNs remember essentials related to the input signal through an internal memory while enabling a prediction of next states.

The main difference of the LSTM is that the gradient can flow for long durations. A crucial modification has been to make the weight on the self-loop conditioned on the context, rather than fixed. The time scale of integration is adjusted by doing the weight of self-loop gated based on the input sequence because the time constants are model output [19]. The architecture of the LSTM is demonstrated in Fig.2.

**Figure 2:**
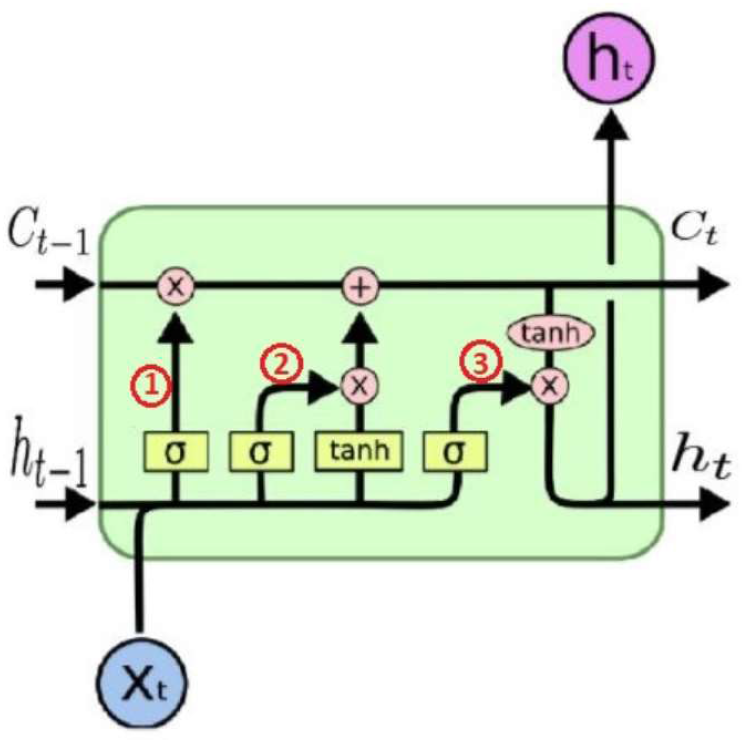
LSTM architecture. 1) Forget gate, 2) Input gate, 3) Output gate

According to Fig.2, × is the scaling of information, + is adding information, *σ* is the sigmoid layer, which is used as a memory for remembering or forgetting, *tanh* is the activation function, which is used to solve the gradient vanishing problem, *h*(*t*) corresponds to the output of LSTM unit, *c*(*t* − 1) denotes the memory from previous LSTM unit, *X*(*t*) is input, *c*(*t*) represents new updated memory. The path from *c*(*t* − 1) to *c*(*t*) is defined as a memory pipeline. Forget gate takes *X*(*t*) and *h*(*t* − 1) as input and decides whether to forget or not the incoming information. Input gate decides what information is stored in memory and output gate choose what information becomes an output.

## 3. MATERIALS AND METHODS

### 3.1. EXPERIMENTAL PARADIGM

BCI Competition IV 2008-Graz data set A provided by the Institute for Knowledge Discovery (Laboratory of Brain-Computer Interfaces, Graz University of Technology), is used in the deep learning-based modelling process. The data set comprises of EEG data from 9 subjects. The BCI paradigm has been established based on the four different motor imagery tasks (**imagination of the movement of the left hand (class 0), right hand (class 1), both feet (class 2), tongue (class 3)**). The session is comprised of **12 runs** having **48 trials** (12 for each of the four existing classes) and are dichotomised by short breaks. **576 trials** are conducted in total.

At first, a recording of approximately 5 minutes was run for estimating EOG interference. The recording is constructed by three-phase [1) two minutes with eyes open, 2) one minute with eyes closed, and 3) one minute with eye movements]. The details of the data acquisition process are represented in [55].

### 3.2. DATA ACQUISITION AND PREPROCESSING

EEG signals were acquired by utilizing twenty-two (22) Ag/AgCl electrodes having inter-electrode distances of 3.5 cm. The montage which maps the EEG and EOG channels are depicted in Fig3. a and Fig3.b, respectively.

**Figure 3:**
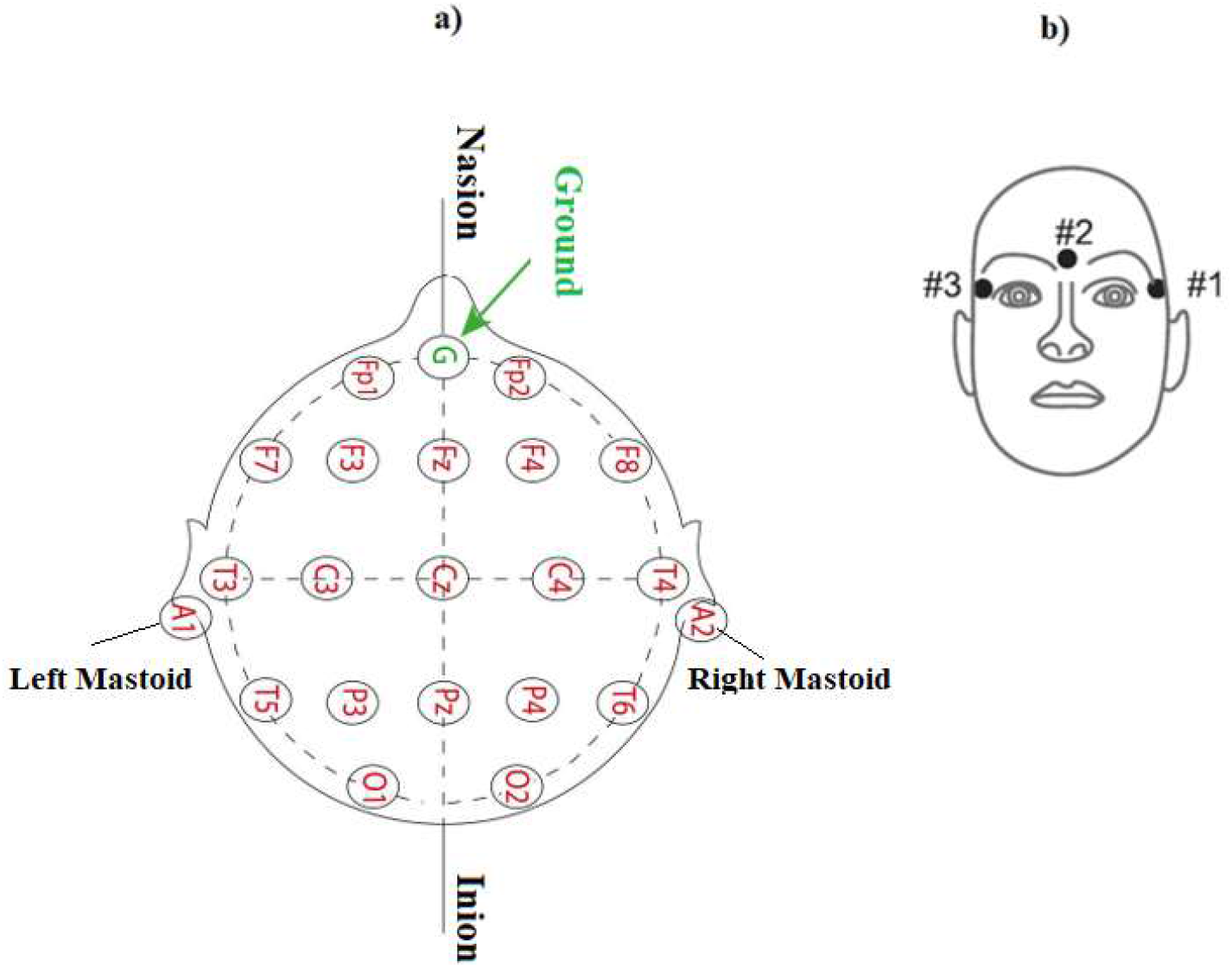
**a)** Electrode montage of the twenty-two (22) channel EEG device. **b)** Electrode montage of the three (3) monopolar EOG channels.

All signals (EEG and EOG) were collected monopolar with the left mastoid as a reference, and the right mastoid as ground. The signal sampling rate is 250 Hz. The signals were filtered with a 0.5-100 Hz. band-pass filter, and with an additional 50 Hz. notch filter, which suppresses the line noise. The sensitivity of the EEG amplifier was set to 100 *μV*. The sensitivity of the EOG amplifier was set to 1mV. The details about the data file description may be accessed from [7]. The data collected from 9 subjects for each mental task was cropped to a total of 196500 sample to equalize the length of the data after NaN values are cleaned. The total size of the processed data is reduced to [196500*4 (sample), 25 (channel-EEG+EOG)] matrix form. After obtaining the final matrix form, the data is standardized by removing the mean and scaling to unit variance. Through the centring and scaling process, the relevant statistics are computed on the samples in the training set. Mean and standard deviation are then stored to be utilized with the transformation. The standardization process should be applied before classification for enhancing the performance of the machine learning-based estimator. They might give poor resuşts when the individual features do not more or less look like standard normally distributed data (e.g. Gaussian with zero mean and unit variance) [12].

### 3.3. ARTIFACT REMOVAL AND DIMENSIONALITY REDUCTION

The signal and noise (both random and deterministic) should be separated to obtain a high-quality measurement. Random noises can be eliminated by repeating a single signal measurement process. Some artefacts originate from the measurement process itself, therefore a statistical analysis of sensors is necessitated [29]. The characteristics of the artefacts differ from the signal of interest. When the artefact is limited to a specific frequency range, it can be removed by a frequency filtering approach. If it is based on discrete frequencies or their harmonics, it can be eliminated by notch filtering. When it is limited to a certain time range, e.g. in the case of eye blinks, the time intervals are discarded by observing the signal. However, if the artefacts are originated from various sources or a limited to a subspace of the signal space, the topography of the artefacts exhibit superposition state. For this reason, the artefacts can be eliminated by utilizing signal-space projection [57], so that the remaining signals do not include artefact subspace [50]. Methods, i.e. ICA [61] or PCA, based on the assumption that artefacts and signal sources are sufficiently independent of each other are also proposed [27; 41]. It is also possible to define the artefact by a particular temporal pattern, i.e. exponential decay. The artefacts are also modelled through fitting its parameters to the data, and then it is removed from the physiological signal, i.e. frequency range, amplitude, etc. The other methods include regression techniques, FIR filters, wavelet denoising, denoising with multilevel wavelet DWT functions, etc. Several artefact removal methods can also be united and run as stand-alone or EEGLAB plug-in. [45]. The suitability of artefact rejection method depends on not distorting the main component or feature sought in the signal. This is especially the case for some automated methods such as (independent or principle) component analysis and some filtering methods. The spatiotemporal nature and the generation mechanisms of the artefacts should be investigated. It is possible to reduce the effects of the muscular and eye-movement artefacts due to their structural difference from simultaneous reference signals such as EOG, ECG, EMG etc. When the large transient artefacts, e.g. from electric stimulation are expected, the recording epoch should be wide enough for confining artefacts in time [56; 42]. The multi-class classification performance of the BCI has also been improved with the current source density (CSD) method, which depends only on the position of the sensors on the scalp [49].

In light of the above basic information, in this paper, the PCA method was preferred to remove artefacts. PCA serves the speed-boosting of the fitting of the classifier by dimensionality reduction. PCA converts data linearly into new features that are not correlated with each other by doing the orthogonal transformation [34].

### 3.4. CNN-LSTM FRAMEWORK

The classifier utilized in this paper is the combination of CNN and LSTM. First, the time-domain features of the EEG data are extracted through 1D-CNN, and after that, these features are feeding into the LSTM to obtain high-level representative features. Finally, the classifier dichotomizes four MI tasks. The framework of the methodology is depicted in Fig.4.

**Figure 4:**
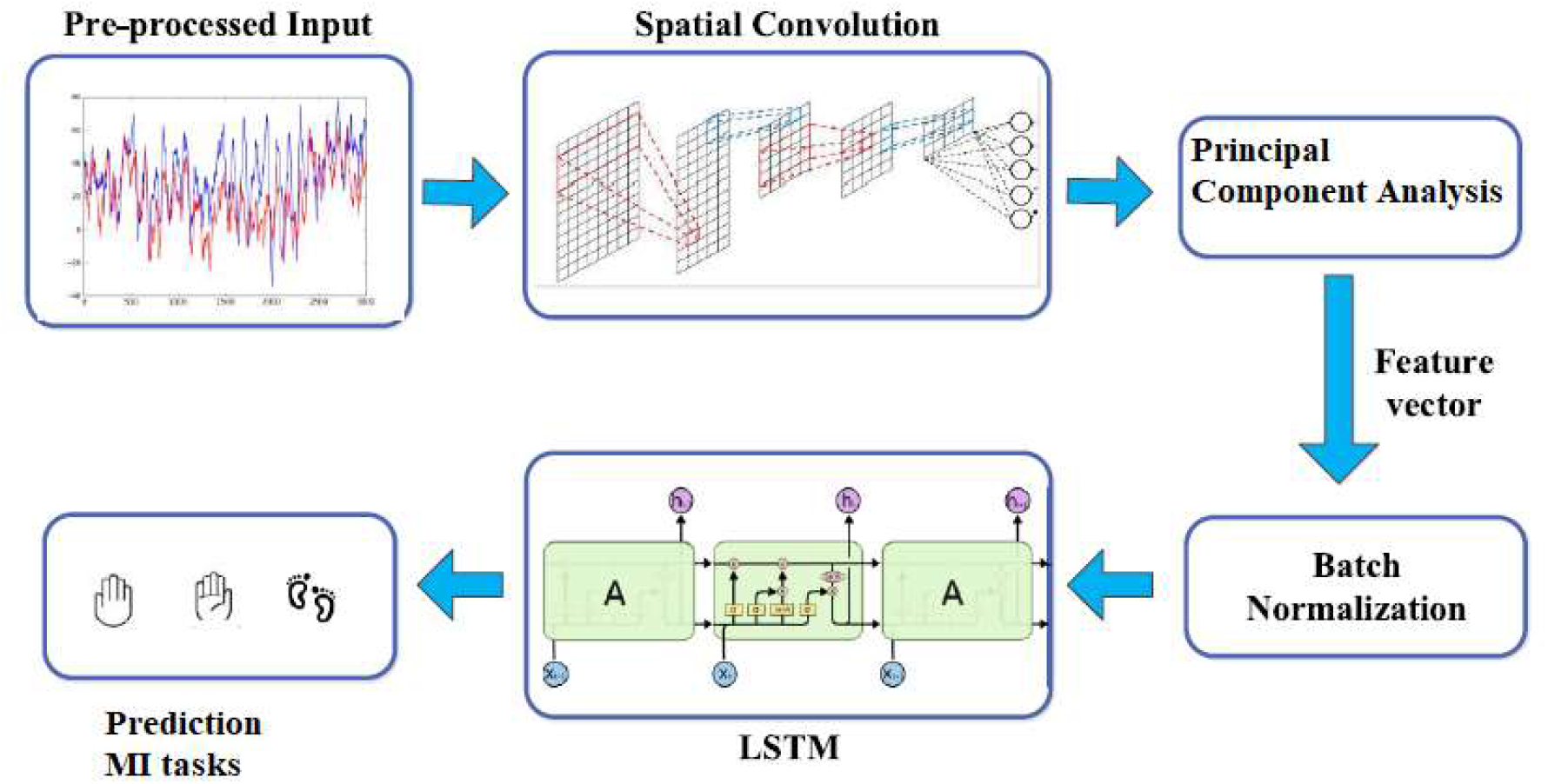
The framework of the classification methodology.

ERD and ERS can be seen in *μ* − *band* (8 − 13 *Hz*.) and *β* − *band* (13 − 30 *Hz*.). For this reason, the *μ* and *β* bands are extracted through the fifth-order butter-worth filter. In this way, it is possible to benefit from band power optimally. In the experiment, the subjects performed MI tasks in 3 seconds, the time segment of 2-6 seconds are extracted to reduce the temporal redundancy.

Multichannel EEG data is two-dimensional, but time and channel have different units, which drives a non-trivial selection of the filter kernel dimensions. The model is based on a hybrid CNN-LSTM structure and the summary of the model is given in Table 1.

**Table 1:**
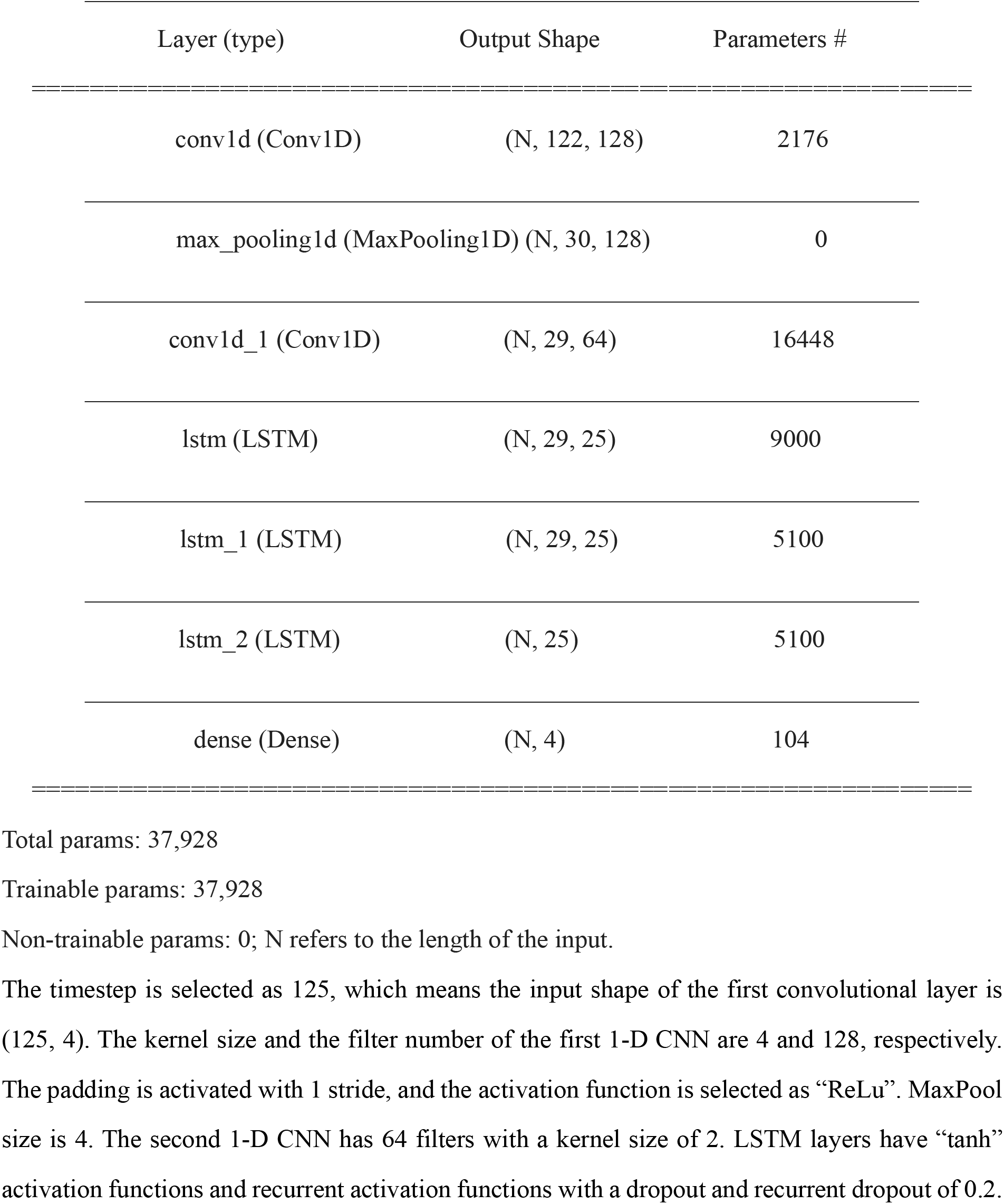
Model summary of the CNN-LSTM. Model: “CNN-LSTM”

The total unit of each LSTM layer is 25. The dense layer has “softmax” activation function. The hyperparameters of the model are represented in Table 2.

**Table 2:** The hyperparameters of the model.

| parameter | type or value |
| --- | --- |
| optimizer | Adam |
| activation | ReLu, tanh, softmax |
| regularization | dropout |
| loss function | categorical cross-entropy |
| batch size | 64 |
| epoch | 400 |

Dropout, which a determined portion of the neuron is randomly turned off at each iteration, is necessary for model generalization. Dataset was shuffled at each epoch to avoid overfitting.

## 4. RESULTS

All steps are executed by the publicly available *“Google Colaboratory”*, which is a free cloud service giving an opportunity to AI developers to apply their deep learning-based algorithms. Datasets are trained on “Google Colaboratory” including several significant modifications, which allows evaluations on multiple GPUs. The GPU model is Tesla k80 supporting Python environment and Keras deep learning libraries. It is easy to upload data from “Google Drive Application” to train the model. Multi-GPU training exploits data parallelism and is carried out by splitting each batch of training images into several GPU batches, processed in parallel on each GPU. The gradient of the full batch is obtained by averaging the computed GPU batch gradients. Gradient computation is synchronous across the GPUs, so the result is precisely the same as when training on a single GPU. First, the filtered datasets are uploaded to “Google Colaboratory”. The size of the dataset matrix is [195000*4 (time-series representing each mental task), 25 (EOG+EEG channels]. Then, the dataset is normalized via StandardScaler command in the sklearn. preprocessing library, and is subjected to the PCA. After the PCA process, the electrode sources are reduced into four main principal components. The obtained size of the dataset matrix becomes [195000*4, 4 (principal component)]. After that, the dataset matrix is divided into timesteps each part includes 125 samples and reshaped as [6288, 125, 4]. Finally, Datasets are split randomly into the train part and test part, for the fitting via sklearn.model_selection.train_test_split command with parameters of test size 0.1, and random state 0.2. After splitting process, the size of the train and test data matrices are [5659, 125, 4], [629, 125, 4], respectively. The CNN-LSTM model is trained with the 10-fold CV method by using train data. This procedure was done by splitting the training dataset into 10 subsets and takes turns training models on all subjects except one which is held out, and computing model performance on the held-out validation dataset. In this paper, 10 models are build and evaluated for CV [37]. For each trial, a sliding window of size 125 along the time axis. The models obtained from the training phase are tested section-by-section, and finally, the average validated the accuracy and kappa value obtained for each section having epochs=400 and batch_size=64.

The results of the 10-fold CV process are represented in Table 3.

**Table 3:**
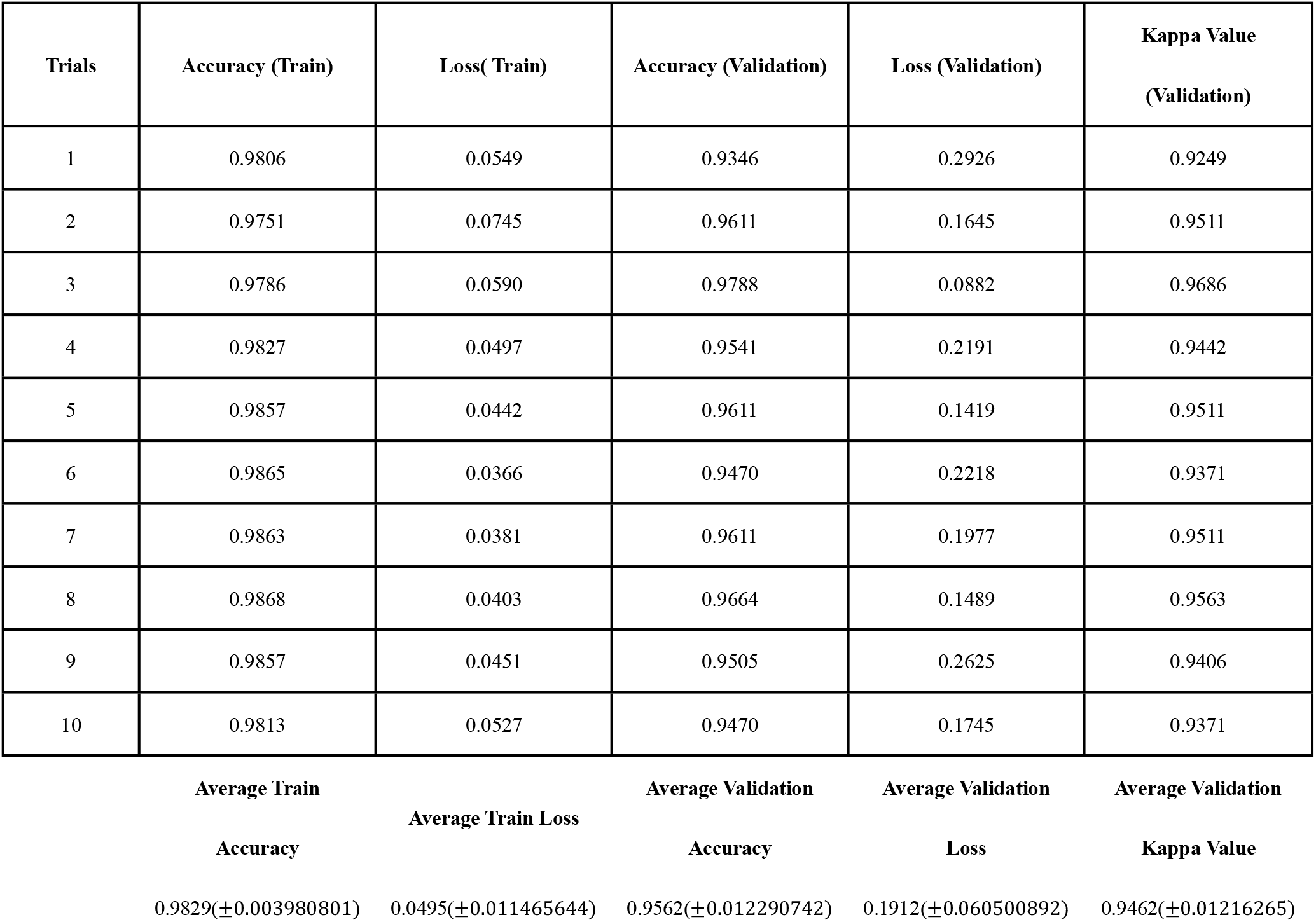
Performance table of the 10-k CV process.

| **Trials** | **Accuracy (Train)** | **Loss( Train)** | **Accuracy (Validation)** | **Loss (Validation)** | **Kappa Value (Validation)** |
| --- | --- | --- | --- | --- | --- |
| 1 | 0.9806 | 0.0549 | 0.9346 | 0.2926 | 0.9249 |
| 2 | 0.9751 | 0.0745 | 0.9611 | 0.1645 | 0.9511 |
| 3 | 0.9786 | 0.0590 | 0.9788 | 0.0882 | 0.9686 |
| 4 | 0.9827 | 0.0497 | 0.9541 | 0.2191 | 0.9442 |
| 5 | 0.9857 | 0.0442 | 0.9611 | 0.1419 | 0.9511 |
| 6 | 0.9865 | 0.0366 | 0.9470 | 0.2218 | 0.9371 |
| 7 | 0.9863 | 0.0381 | 0.9611 | 0.1977 | 0.9511 |
| 8 | 0.9868 | 0.0403 | 0.9664 | 0.1489 | 0.9563 |
| 9 | 0.9857 | 0.0451 | 0.9505 | 0.2625 | 0.9406 |
| 10 | 0.9813 | 0.0527 | 0.9470 | 0.1745 | 0.9371 |

|  |  |  |  |  |
| --- | --- | --- | --- | --- |
| **Average Train** | **Average Train Loss** | **Average Validation** | **Average Validation** | **Average Validation** |
| **Accuracy** |  | **Accuracy** | **Loss** | **Kappa Value** |
| 0.9829($\pm 0.003980801$) | 0.0495($\pm 0.011465644$) | 0.9562($\pm 0.012290742$) | 0.1912($\pm 0.060500892$) | 0.9462($\pm 0.01216265$) |

The mean and standard values of the validation accuracy and validation kappa value after 10-fold CV are evaluated as 95.62 % (±1.2290742) and 0.9462 (±0.01216265), respectively. After that, the CNN-LSTM model was tested by using test data. The accuracy and kappa value were obtained as 96.98 % and 0.9597, respectively. The disadvantage of the k-fold CV is that the size of the train test splits is predetermined. The metric of the kappa score evaluation is *“cohen_kappa_score”*, which is a statistic that measures inter-annotator agreement. The details and theoretical basis are explained in [9; 3]. The metrics are implemented to measure classification performance by sklearn.metrics module.

The model accuracy and model loss values of test and train data for each epoch was represented in Fig.5a and Fig.5b, respectively.

**Figure 5:**
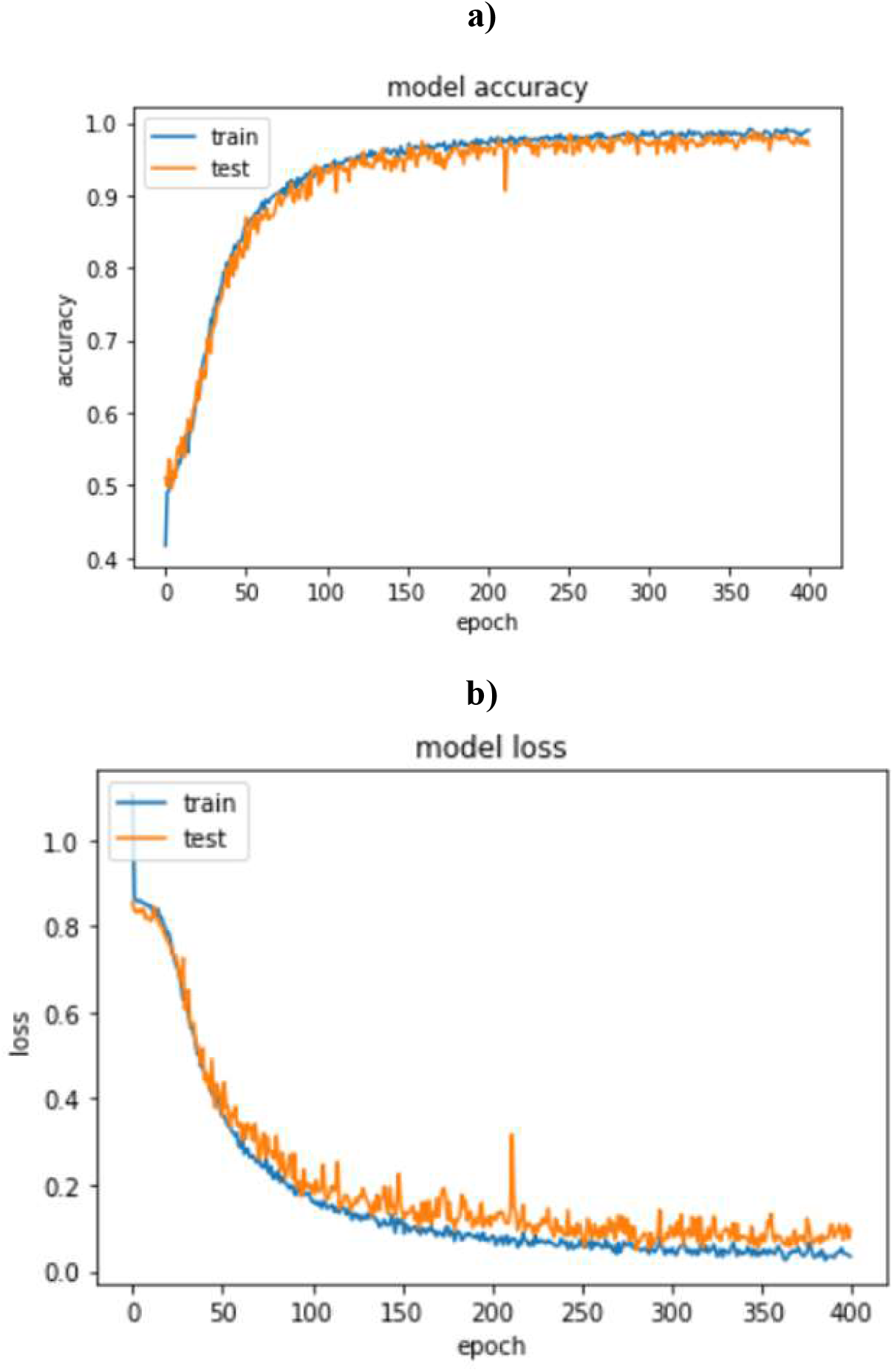
The performance metrics of the model obtained from test and train data a) Model accuracy b) Model loss

The classification accuracy is computed from the confusion matrix with each row corresponding to the true class. In Fig.6, the test confusion matrix was plotted.

**Figure 6:**
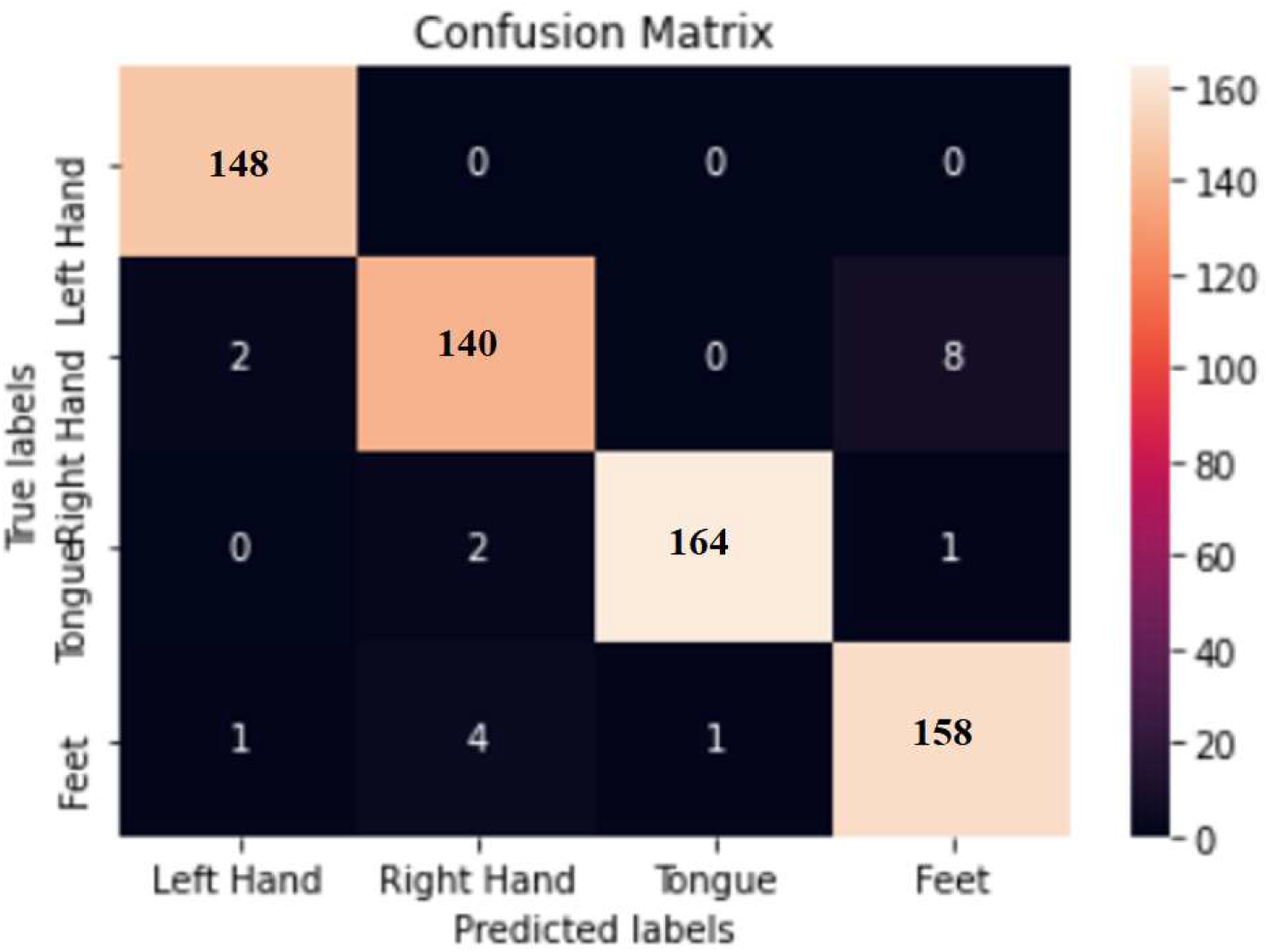
The confusion matrix representing each MI class obtained from test data.

According to Fig.6, the diagonal elements demonstrate the number of points for which the predicted label is equal to the true label, while off-diagonal elements are those that are mislabeled by the classifier. The higher diagonal values of the confusion matrix the better, showing many correct predictions.

When target classes are not balanced, the accuracy metric may not be the right measure. Therefore, the additional metrics like Precision, Recall, F Score etc., should be considered. In Table 4, the results corresponding to these metrics are tabulated.

**Table 4:** Additional metrics showing the classifier performance.

|  | *precision* | *recall* | *f1-score* | *support* |
| --- | --- | --- | --- | --- |
| *Left Hand* | 0.98 | 1.00 | 0.99 | 148 |
| *Right Hand* | 0.96 | 0.93 | 0.95 | 150 |
| *Tongue* | 0.99 | 0.98 | 0.99 | 167 |
| *Feet* | 0.95 | 0.96 | 0.95 | 164 |
| *macro avg* | 0.97 | 0.97 | 0.97 | 629 |
| *weighted avg* | 0.97 | 0.97 | 0.97 | 629 |

A macro-average compute the metric independently for each class and then take the average, whereas a micro-average aggregate the contributions of all classes to evaluate the average metric. The ROC curve was also plotted in Fig.7. The ROC curve indicates the “True Positive Rate” (Sensitivity/Recall) against the “False Positive Rate” (1-Specificity) at various classification thresholds.

**Figure 7:**
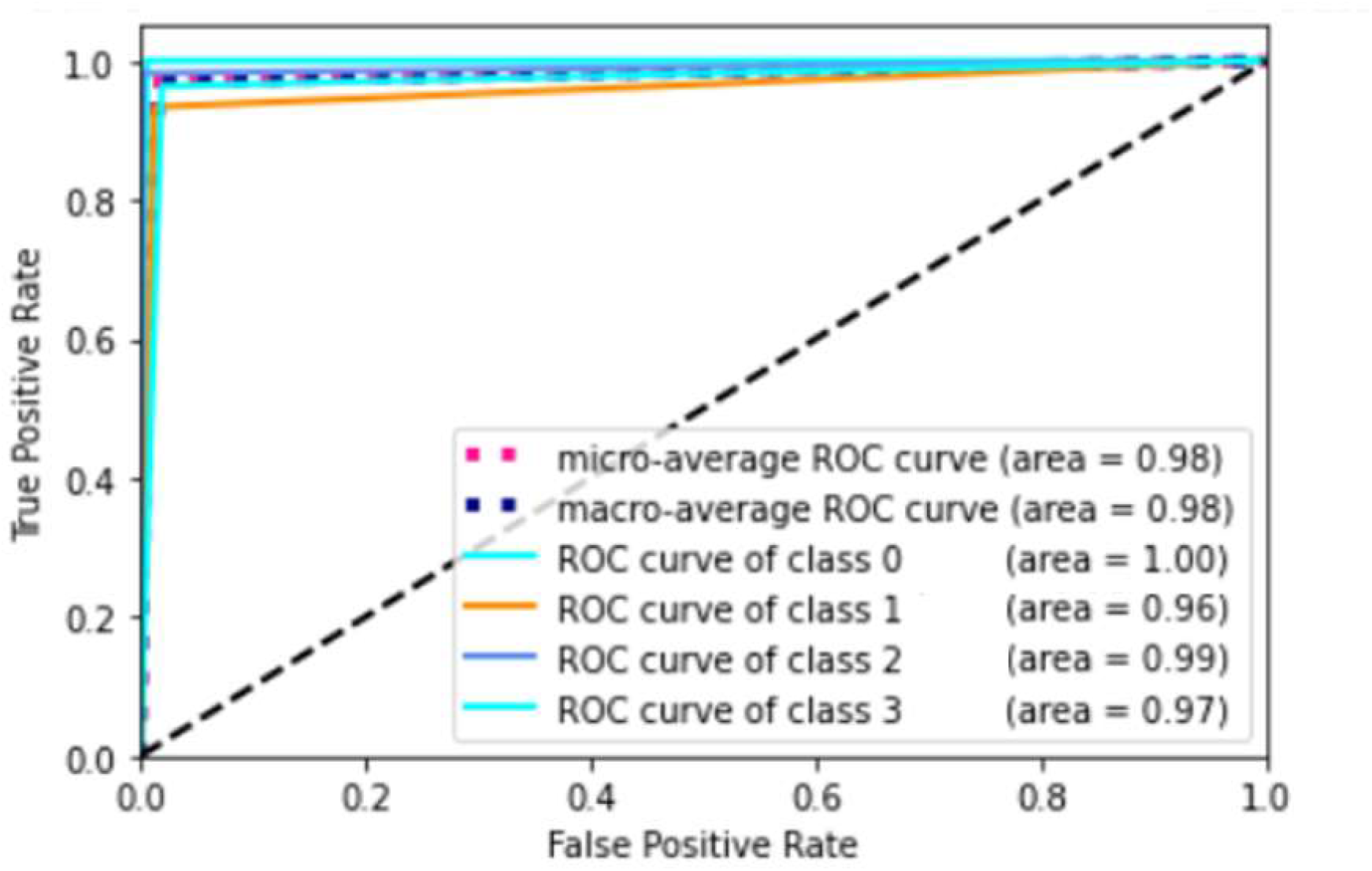
ROC curve. Class 0: Left Hand, Class 1: Right Hand, Class 2: Tongue, Class 3: Feet

ROC curve is adapted to multi-label classification case by binarizing the output. One ROC curve can be drawn per label, but one can also draw a ROC curve by considering each element of the label indicator matrix as a binary prediction (micro-averaging).

AUC measures the entire two-dimensional area underneath the curve. How well a parameter can distinguish among two diagnostic groups can be measured by evaluating the AUC score. The AUC score is estimated as 0.9798126815598498. AUC provides an aggregate measure of performance across all possible classification thresholds. Interpreting AUC is as the probability that the model ranks a random positive example more highly than a random negative example. AUC is scale-invariant and classification-threshold-invariant [20].

## 5. CONCLUSIVE SUMMARY AND DISCUSSION

The performance of the CNN-LSTM model was evaluated robustly by 10-Fold CV. According to the results, the hybrid CNN-LSTM model achieved a quite satisfactory and reliable accuracy and kappa value. The predictive model has also extracted the most relevant information from the beginning and the end of the imagined movements. The choice of such a recurrent model depends on the requirement of increasing prediction accuracy, assuming that there is never an abruptly change of movement type in the given experiment. The robustness against overfitting was regularized by adding dropout and pooling layers, doing k-fold CV, and making a batch learning process, which averages over 64 samples from various subjects in each training step. This framework can achieve superior performance in MI classification tasks, and the robustness on different subjects can be improved with appropriate filtering and initial weights. The results are comparable with the reported accuracy values in related studies and the designed CNN-LSTM architecture outperforms the results in the literature [55; 6] on the same underlying data given that the model can learn features from data without necessitating a specialized feature extraction methods.

As a result of this study, it is highly recommended to utilize the combination of the CNN-LSTM based DL architecture for building a BCI system. It is worth investigating the possibilities of LSTM for any real-time sequence classification having online feedback.

